# A Mean-Field Approach to Criticality in Spiking Neural Networks for Reservoir Computing

**DOI:** 10.1101/2025.02.05.636716

**Authors:** Ruggero Freddi, Francesco Cicala, Laura Marzetti, Alessio Basti

## Abstract

Reservoir computing is a neural network paradigm for processing temporal data by exploiting the dynamics of a fixed, high-dimensional system, enabling efficient computation with reduced complexity compared to fully trainable recurrent networks. This work presents an analytical framework for configuring in the critical regime a reservoir based on spiking neural networks with a highly general topology. Specifically, we derive and solve a mean-field equation that governs the evolution of the average membrane potential in leaky integrate-and-fire neurons, and provide an approximation for the critical point. This framework reduces the need for an extensive online fine-tuning, offering a streamlined path to near-optimal network performance from the outset. Through extensive simulations, we validate the theoretical predictions by analyzing the network’s spiking dynamics and quantifying its computational capacity using the information-based Lempel-Ziv-Welch complexity near criticality. Finally, we explore self-organized quasi-criticality by implementing a local learning rule for synaptic weights, demonstrating that the network’s dynamics remain close to the theoretical critical point. Beyond AI, our approach and findings also have significant implications for computational neuroscience, providing a principled framework for quantitatively understanding how biological networks leverage criticality for efficient information processing.

## Main

Reservoir computing (RC) is a computational framework tailored for efficiently processing temporal data by leveraging the inherent dynamics of a largely fixed reservoir, a high-dimensional recurrent dynamical system that transforms input signals into complex internal states (Schuman et al. 2022; Tanaka et al., 2019). This transformation allows a readout layer to learn target outputs with reduced computational overhead compared to fully trainable recurrent neural networks, which require optimization of all connection weights (Werbos, 1999). RC has demonstrated success in diverse applications, including machine vision, time series forecasting, classification, anomaly detection, and system identification (Tan and van Dijke, 2023; Kärkkäinen and Linna, 2022; see also Tanaka et al. 2022, and the references therein).

Reservoir architectures vary widely in implementation. One prominent RC model, the Echo State Network (ESN; Jaeger, 2001; Jaeger and Haas, 2004), employs a sparsely connected network of hidden neurons with fixed random weights. These weights satisfy the echo state property, ensuring that previous states influences decay over time, with the reservoir’s dynamics driven predominantly by input signals. Another key RC variant, the Liquid State Machine (LSM; Maass et al., 2002), utilizes a spiking neural network (SNN) as its reservoir, thus drawing inspiration from cortical dynamics in biological brains.

The choice of neuron model is critical for reservoir design, as it directly impacts computational performance and functional capacity. In the realm of those who opt for biologically realistic models, there exists a vast complexity spectrum of possibilities. At the one end of this spectrum lie the detailed models, such as the Hodgkin-Huxley model (Hille, 2001; Hodgkin and Huxley, 1952). These models offer precise descriptions of biological neural activity but are computationally intensive, making them well-suited for capturing the behavior of a single neuron, yet impractical for large-scale artificial neural network applications.

In such a context, simplified neuron models like the leaky integrate-and-fire (LIF) neuron are commonly preferred. LIF neurons balance computational efficiency with the ability to reproduce essential neural features, such as spike-based temporal integration (Teeter et al., 2018). These lightweight models can be extended to incorporate additional features of interest and are well-suited for both ESNs and LSMs (Ebato et al., 2024; Jaeger et al., 2007), resulting in almost identical definitions. The LIF model captures the fundamental functional behavior of neurons without the computational demands of detailed ion-channel modeling, thus serving as a practical choice for spiking reservoirs designed to process spatiotemporal data effectively and at scale.

In SNN-based reservoir implementations, it is crucial to position the dynamical system at the edge of chaos, i.e., near a suitable order-disorder phase transition point, as it serves as the primary computational component. Operating in this regime enhances the reservoir’s ability to process temporal data with heightened sensitivity and adaptability, while fostering complex, diverse dynamics and achieving versatile computational performance (Woo et al., 2024; Kinouchi and Copelli, 2006; Langton, 1990).

Achieving this requires careful tuning of the reservoir hyperparameters. In particular, the average synaptic weight is a pivotal factor in optimizing the regime of an SNN, as it directly affects the inter-spike interval (ISI), which is straightforwardly linked to the network’s firing rate (Adams et al., 2024; Koralalage et al., 2023).

Levina et al. (2007) analyzed the self-organized criticality of an integrate-and-fire model with transmission delays, but without leakage, deriving a relationship between ISI and average synaptic weight. From this, it is theoretically possible to analyze the presence of critical points. Nevertheless, their model excludes refractory periods and, more importantly, ties the relation to the mean avalanche size, which cannot be used as a hyperparameter in the reservoir implementation.

Brochini et al. (2016) analyze a special case of the Galves-Löcherbach model, deriving analytical expressions for the firing rate density at the critical point. Nonetheless, their analysis assumes no memory effects, is limited to fully connected networks, and rely on the assumption of infinite network size, whereas reservoir computing implementations must contend with finite-size effects.

In the absence of a theoretical expression for optimal weight initialization, synaptic weights are often set heuristically and subsequently refined through Hebbian learning rule, such as spike-timing-dependent plasticity (STDP), which adjusts weights based on the precise timing of spikes (Norton and Ventura, 2006). Extensions of these approaches, such as models incorporating astrocytes, have been proposed to enhance network performance and promote operation near excitation-inhibition balanced criticality (Ivanov and Michmizos, 2021), a condition widely recognized as essential for maintaining chaotic network activity and optimizing information processing (Barzon et al., 2024; Brunel, 2000; van Vreeswijk and Sompolinsky, 1996). However, these strategies predominantly focus on computational learning and adaptive tuning rather than exploiting a direct relation to determine initial conditions. A promising alternative would be to explicitly identify the critical point as a function of the mean synaptic weight in the finite-size case and initiate network dynamics directly within this regime. This approach could bypass the need for extensive fine-tuning, allowing the reservoir to operate optimally from the outset.

In this work, we present an analytical framework for configuring SNN-reservoirs in the critical regime by deriving and solving a mean-field equation that describes the evolution of the average membrane potential in a LIF neural network. The model is highly general, accommodating a wide range of network topologies, from sparse to fully connected, where reservoir nodes interact and may receive randomly distributed external inputs. Our analysis establishes a theoretical relation between the average synaptic weight and ISI (2) (Panel A, Fig. 1), yielding an explicit approximation of the critical point (3). Notably, the approximation reduces the latter relation to one equivalent to its leak-free counterpart, for which we provide a complete analytical treatment.

**Fig. 1.**
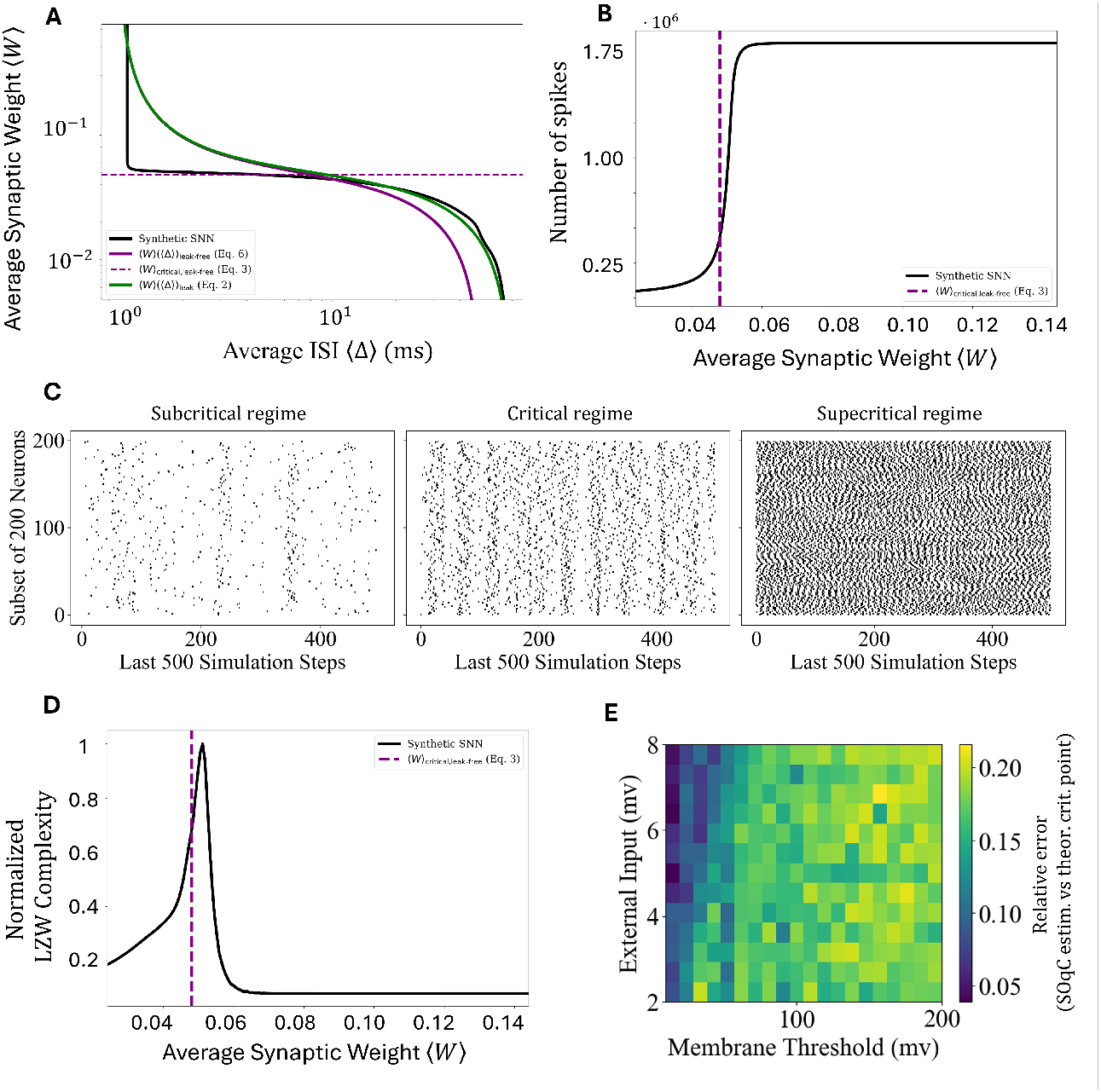
Panel A illustrates the theoretical relation between the average synaptic weight and ISI in the leaky system (green solid curve; (2)) and in the leak-free system (purple solid curve; (6)), alongside the curve estimated from synthetic LIF small-world network data. It is important to emphasize that both the solid lines, representing the median over the iterations, and the shaded areas, indicating the interquartile range, are plotted in the panels. However, the shaded areas are extremely small due to the high stability of the results, making them barely visible. The dotted purple horizontal line represents the estimated critical point, whose expression is given in (3). The hyperparameter values used in the simulations are detailed in the Methods section. Panel B shows how the number of spikes in the SNN varies as a function of the network’s average weight. The estimated critical point (dotted purple vertical line) is close to a drastic change in the dynamics. Panel C displays a raster plot of the activity of 500 SNN neurons during the last 500 time steps of the simulation for three different mean synaptic weights, corresponding to 80%, 100%, and 110% of the critical weight given by our theoretical approximation. The first case exhibits sparse activity, while the last shows highly deterministic oscillatory behavior. The critical regime, positioned between these two extremes, clearly represents a balance between the two activity patterns. Panel D demonstrates that the estimated critical point aligns closely with the peak of information-based complexity, indicating that it represents the most informative state of the system. Panel E presents the absolute relative error between the final synaptic weight of the synthetic SNN and the theoretical value in a simple computational example of self-organized quasi-criticality. Even with a simple learning rule, the system converges toward the theoretical value. However, our analysis enables real-time updates of the critical point by directly leveraging equation (3).

To validate our predictions, we simulate the LIF network’s spiking dynamics across various synaptic weights and confirm that the system exhibits critical behavior near the predicted critical point (Panel B-C, Fig. 1). Additionally, we evaluate the benefits of operating near criticality by computing the Lempel-Ziv-Welch complexity (Welch, 1984) of synthetic spike trains, demonstrating the enhanced computational richness of reservoirs in the critical regime (Panel D, Fig. 1). Finally, we implement a toy model of self-organized quasi-criticality (SOqC) through a local learning rule for synaptic weights, showing that the dynamics remain close to the theoretically predicted critical regime (Panel E, Fig. 1).

## Results

We consider a reservoir composed of a network of *N* leaky integrate-and-fire neurons. The *i*-th neuron emits a spike at time *t* if its membrane potential *v*_*i*_(*t*) exceeds a threshold *θ >* 0. Upon spiking, the membrane potential resets to *v*_reset_ = 0, and the neuron enters a non-negative refractory period *τ*_ref_. The evolution of the membrane potential is governed by three main components. The first component accounts for the fact that the network is partitioned into *K* ≤ *N* groups, with the *k*-th subset composed of *n*_*k*_ neurons. In this setup, every *τ >* 0 seconds, one neuron is uniformly selected. If the selected neuron (e.g. the *i*-th) belongs to the *k*-th group, it receives an external input that alters the potential proportionally to a resistance *R*_*k*_, such that 0 *< R*_*k*_*I*_*k*_ *< θ*. It is worth to notice that the partition of neurons into subset associated with different external currents reflects the practical design of reservoirs in which only certain neurons are directly connected to the input nodes. In such cases, the external currents for neurons not receiving input are set to zero, ensuring that only the designated input-connected neurons collect the external signals. The second component accounts for the cumulative effect of synaptic inputs from other *C*_*i*_ neurons connected to *i*. Each synapse contributes with a discrete spike of weight *W*_*i*,*j*_ (which is equal to 0 in the case of no connection) to *v*_*i*_(*t*) whenever neuron *j* fires, with the timing determined by 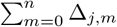, where Δ_*j*,*m*_ is the inter-spike interval for neuron *j* at its *m*-th firing. This component aggregates the contributions of all valid presynaptic neurons up to time *t*. The final component represents an exponential decay in the membrane potential at a rate *α*. This term models the natural tendency of the neuron to leak charge, causing *v* to decay over time in the absence of input.

Accordingly, the dynamics of the membrane potential for neuron *i* in the *k*-th group is governed by the following differential equation:

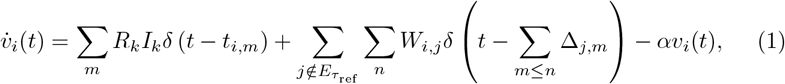

where *t*_*i*,*m*_ denotes the time at which neuron *i* receives its *m*-th external input, and 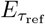 the set of neurons in refractory period.

Using a mean-field approximation, we analyze the evolution of the average contribution ⟨*v*(*t*)⟩_*k*_ for the *k*-th group and, subsequently, the overall average membrane potential ⟨*v*(*t*)⟩ := Σ _*k≤K*_ *n*_*k*_⟨*v*(*t*)⟩_*k*_*/N* . The mean-field formulation approximates the inter-spike intervals Δ_*j*,*m*_, the synaptic weights *W*_*i*,*j*_, and neuron degree with their average values, ⟨Δ⟩, ⟨*W* ⟩, and *βN* with *β* ∈ (0, 1), respectively. From the solution ⟨*v*(*t*)⟩, assuming ⟨*v*(0)⟩ = *v*_*reset*_ = 0 and terming *RI* := Σ_*h*_ *n*_*h*_*R*_*h*_*I*_*h*_*/N*, we derive the functional relationship between the average synaptic weight ⟨*W* ⟩ and the mean inter-spike interval ⟨Δ⟩, given by:

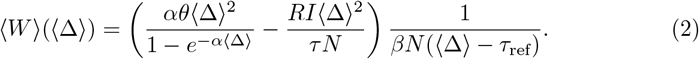

We consider the conditions *ατ*_ref_ *<* 1*/*2, *θτ N/*(*RIτ*_ref_) ≥ 2, and *θτ Nα/*(*RI*) *<* 1, which, as detailed in the Methods section, ensure the theoretical results remain physically plausible and validate our analysis of criticality. A graphical representation of the relationship (2) alongside experimental observations is provided in Panel A of Fig. 1.

In our framework, a critical point corresponds to a state where a thermodynamic function undergoes a drastic change, signaling a phase transition in the system. At such points, the function itself or its higher-order derivatives diverge or become discontinuous. Specifically, under a relaxed hypothesis, we aim to approximate the expression of the point that causes the derivative of ⟨*W* ⟩ (⟨ Δ ⟩) to approach zero. This corresponds to the condition where the inverse function ⟨Δ⟩ (⟨ *W* ⟩) exhibits a nearly diverging first derivative. To achieve this, we analyze a related system that serves as an approximation. In particular, (2), under the validity of the three above defined conditions, can be approximated with a function that we have proved to characterize the leak-free system under the mean-field assumptions.

For the leak-free system, we derive the exact theoretical relationship between the ISI and the average synaptic weight (the inverse of the approximation of (2)), as shown in (7). Furthermore, we analytically determine the critical ISI, as presented in (8), along with the corresponding critical average synaptic weight ⟨*W* ⟩_critical,leak-free_. The latter quantity may correspond to an approximation of the critical point of the original system. That is, denoting the critical average synaptic weight of the LIF network of interest as ⟨*W* ⟩_critical_,

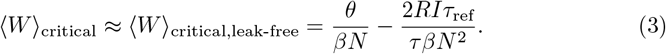

The results offer a bird’s-eye view of how the hyperparameters can change the position of the criticality in the (⟨*W* ⟩, ⟨Δ⟩) plane, enabling the possibility of a top-down modulation of the system to position it at the desired operating point. It is worth noting that, when *βN* is large, the expression of the critical point corresponds to a relatively small value, which aligns with the standard concept of the balance between excitation and inhibition. This observation highlights the delicate interplay necessary to maintain stability within the system, ensuring that neither excitation nor inhibition dominates excessively.

To computationally validate the analyzed critical regime in case of LIF system, we rely on simulations of the SNN reservoir dynamics. Panel B of Figure 1 depicts the number of spikes as a function of ⟨*W* ⟩ (hyperparameters’s configuration detailed in the Methods section). The spike count graph demonstrates a drastic change close to the critical point estimated using (3), consistent with the occurrence of a phase transition in the network. In the subcritical regime (⟨*W* ⟩ *<* ⟨*W* ⟩ _critical_), the number of spikes is low, reflecting a rare and sparse firing activity indicative of a quiescent state where activity is prone to extinction. Conversely, in the supercritical regime (⟨*W* ⟩ *>* ⟨*W* ⟩ _critical_), the spike count sharply increases, approaching values consistent with the maximum number that the network can generate. This neuron’s behavior aligns with a regime characterized by oscillatory activity, alternating between periods of firing and silence. These observations reinforce the notion that the critical point marks a fundamental transition in the network’s dynamics, separating a low-activity regime from a highly active and oscillatory state (see Panel C of Fig. 1 for a raster plot of the SNN’s activity in the three different regimes).

Panel D of Fig. 1 illustrates the normalised information-based Lempel-Ziv-Welch (LZW) complexity, as a function of the average synaptic weight, under the same conditions as the previous simulation. The complexity exhibits a pronounced peak close to the theoretical critical point. Across the different experimental configurations tested (chosen to evaluate the results under potentially challenging conditions, such as the boundaries of the validity domain), the absolute relative errors between the peak of the complexity curve and the critical point remained below 0.26, with 60% of the tested configurations showing an error smaller than or equal to the algorithm’s precision. Nonzero errors consistently corresponded to an underestimation of the actual peak by the value in (3). The error may be mitigated e.g. by increasing the simulation duration or by moving away from the validity boundaries of the conditions.

The presence of a clear peak in complexity highlights the fundamental transition in the network’s dynamics. In the subcritical phase, the complexity remains low, reflecting the network’s behavior of extinguishing activity. Instead, in the supercritical phase, the neuron’s state almost deterministically alternates between firing and being quiescent. This deterministic alternation can be attributed to the interplay of the refractory period and the dynamics of synaptic interactions. This limits the achievable LZW complexity compared e.g. to a Bernoulli process with *p* = 0.5, where each state (1 or 0) is equally probable and independent of prior state. This analysis underscores that the region around the critical point is the most relevant in terms of information content and computational capability. It is worth noting that the results described above can also be achieved in networks with as few as 100 neurons, demonstrating the approach’s robustness and suitability for low-power devices with strict hardware constraints.

Finally, we showcase a computational example illustrating the SNN’s ability to self-organize near the theoretical critical point via a simple local learning rule, which aims to converge toward a state where only half of the neurons fire within a given *τ*_ref_ period. In particular, for all analyzed configurations, the absolute relative errors between the mean weight at which the dynamics settle and the critical point in (3) are observed to lie within the range 0-0.3 (Panel E of Fig. 1). We observed that during the progression of the simulation, the results remain close to the critical point once it is approached, indicating that from that point onward, they no longer significantly depend on the experiment length. Nevertheless, it is worth noting that by exploiting (3), the system can be initialized in a near-optimal state, reducing reliance on such rules or, at the very least, simplifying their convergence toward this state. Furthermore, any temporal variation in hyperparameters, such as input intensity, which inevitably causes a slight shift in the critical point, can be addressed by directly updating the mean synaptic weight using our theoretical findings.

## Discussion

We propose a detailed framework for implementing a spiking neural network (SNN) reservoir designed to operate near a critical regime from the outset. This can be achieved by precisely tuning the average synaptic weight to match the analytically derived expression of the critical point of an approximated network model. To obtain this expression, we conducted an in-depth analysis of leaky integrate-and-fire networks with highly general topologies under the mean-field approximation. We also provided a rigorous analytical treatment of integrate-and-fire SNN and validated the theoretical results through synthetic experiments. Such experiments included Several hyperparameter settings, e.g., topology ranging from sparse to fully connected and from randomly uniform to small-world structure.

Our findings extend previous research on criticality and synaptic weight balance (e.g., excitation/inhibition) by addressing cases with a finite number of neurons, as encountered in SNN-reservoir implementations. Additionally, we exclusively utilize hyperparameters that are inherently suited for defining the network, e.g. the neurons’ average connection degree, without requiring them to diverge or to approach zero. Our results shed light on how these external configuration variables influence the position of the edge of chaos, uncovering the coupling between hyperparameters and fundamental network properties, such as the generated information content and the system’s activity level.

Simulations reveal that critical behavior can be reliably achieved in networks with as few as 100 neurons, demonstrating that our proposed approach maintains robustness and scalability even in resource-constrained scenarios. This highlights its feasibility for deployment in low-power computational devices, such as some portable systems, where hardware limitations impose strict constraints on network size and energy efficiency.

Finally, our results provide deeper insights into the functioning of biological neural networks, offering a theoretical framework that bridges computational neuroscience and practical AI applications. This lays the foundation for further exploration of self-organization and the analysis of criticality in more complex neural models for reservoir computing, such as nonlinear extensions of LIF networks.

## Methods

### Mean-field approximation

Under the mean-field approximation, the first term of (1) primarily consists of impulses arriving at a frequency of *n*_*k*_*/*(*τ N*)(1*/n*_*k*_) = 1*/*(*τ N*), as each neuron is selected with uniform probability. In the second term, the local synaptic weights *W*_*i*,*j*_ for the *k*-th group can be approximated by the average weight ⟨*W* ⟩_*k*_, and the inter-spike intervals Δ_*j*,*m*_ by their average value across the whole network, i.e. ⟨Δ ⟩ ; the average instantaneous contribution of presynaptic spikes to the membrane potential, assuming that neurons are connected to a portion of the total population, i.e., Σ _*i*_ *C*_*i*_*/N* = *βN* with *β* ∈ (0, 1), and that only a fraction of them are available to emit spikes at a given moment, i.e. 1 − *τ*_ref_*/* ⟨Δ ⟩, is then determined as ⟨*W* ⟩_*k*_*βN* (1 − *τ*_ref_*/* ⟨Δ ⟩)*/* ⟨Δ ⟩. Hence, averaging across the *K* subsets,

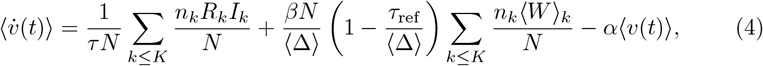

whose solution, assuming ⟨*v*(0)⟩ = *v*_reset_ = 0, terming *RI* := Σ_*k≤K*_ *n*_*k*_*I*_*k*_*/N*, and ⟨*W* ⟩ := Σ _*k*≤*K*_ *n*_*k*_⟨*W* ⟩_*k*_*/N*, is

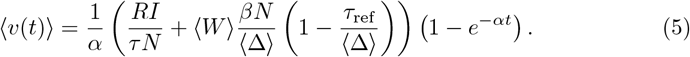

Assuming that the ISI associated with ⟨*v*(*t*)⟩ corresponds to ⟨Δ⟩, the average membrane potential is periodic, satisfying ⟨*v*(⟨Δ⟩)⟩ := lim_*t*→⟨Δ⟩ −_ ⟨*v*(*t*)⟩ = *θ*. By exploiting the latter equality, for ⟨Δ⟩ ≥ *τ*_ref_, we have the equation (2). We can now analyze this quantitative relationship in two regimes: one where ⟨Δ⟩ tends to 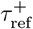, and the other where ⟨Δ⟩ tends to +∞.

The conditions 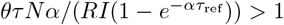 and *θτ Nα/*(*RI*) *<* 1 ensure that the function diverges to +∞ as ⟨Δ⟩ approaches 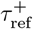, and diverges to −∞ as ⟨Δ⟩ approaches +∞. The validity of these behaviors is crucial, as it is expected that for large positive mean weights, the system fires as rapidly as possible limited by ⟨Δ⟩ ≥*τ*_ref_. Conversely, a large negative mean weight prevents the system from generating any spike, resulting in an infinite ISI. Moreover, the aforementioned constraints ensure that the leak term does not render the temporal distribution of spikes significant, thereby preserving the validity of the mean-field approach. They also guarantee that the current supplied to the system during the refractory period does not lead to saturation. In the next section, the first condition will be replaced to account for the existence of the system’s critical point under consideration.

### Critical Point Estimation

For *α* ⟨Δ⟩ ≪ 1, the term (1 − *e*^−*α*⟨Δ⟩^) behaves like *α* ⟨Δ⟩. As a result, the system behaves similarly to the one without the leak. By deriving (2) from (4) under the assumption of a leak-free model (i.e., by setting the last term of (4) to zero, solving the resulting differential equation, and expressing the average synaptic weight as a function of the average inter-spike interval), we obtain an expression that is identical to (2) when the term (1 − *e*^−*α*⟨Δ⟩^) is substituted with *α*⟨Δ⟩, i.e.,

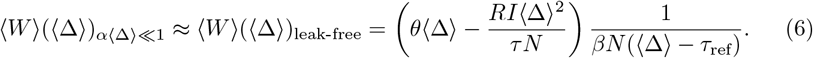

Our goal is to determine the point where the slope of the derivative of the inverse function of (7) on its valid branch, i.e.,

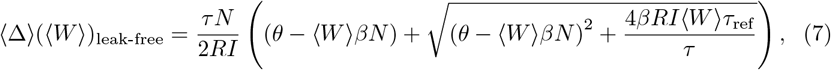

is at its steepest. To achieve this, we analyze its behavior by examining the second derivative of (7).

We observe that the numerator of the second derivative is equal to 8*RIβ*^2^*τ*_ref_*/τ* (*θ* − *RIτ*_ref_*/*(*τ N*)), while the denominator, which is always positive, is given by ((*θ* − ⟨*W* ⟩*βN*)^2^ + (4*RI*⟨*W* ⟩*βNτ*_ref_*/*(*τ*))^3/2^ . Thus, the second derivative is positive, and consequently the first derivative is increasing, if and only if *θτ N/*(*RIτ*_ref_) *>* 1. Since the limit of the first derivative as ⟨*W*⟩ → + ∞ approaches 0, when it is increasing, it is also always negative, meaning that the function in (7) is decreasing. Based on the discussion above, the second derivative achieves its maximum at the value shown in (3). Accordingly, this represents the critical point under the relaxed condition of large but finite value, and the corresponding critical value equals

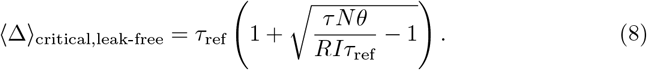

It is important to note that the value in (3) remains non-negative when *θτ N/*(*RIτ*_ref_) ≥ 2. Since *θτ N/*(2*RIτ*_ref_) *> θτ Nα/*(*RI*) for *ατ*_ref_ *<* 1*/*2, and 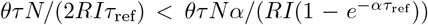, the conditions to be satisfied, replacing those described in the previous subsection, are *ατ*_ref_ *<* 1*/*2, *θτ N/*(*RIτ*_ref_) ≥ 2, and *θτ Nα/*(*RI*) *<* 1. These conditions also ensure the validity of the approximation *α*⟨Δ⟩ ≪ 1.

### Computational validation of criticality

We provide an investigation of the network’s spiking dynamics (in terms of the number of spikes and the information-based Lempel-Ziv-Welch complexity; Welch, 1984) as a function of the average synaptic weight; this aims to computationally validate the critical regime in which the reservoir may directly operate when the theoretical relationship (2) is exploited.

The SNNs consist of directed small-world networks of *N* = 1000 LIF neurons, each firing when its membrane potential exceeds a threshold *θ* = 5 mV and entering a refractory period *τ*_ref_ = 1 ms. Synaptic weights are initialized from a Gaussian distribution 𝒩 (⟨*W*⟩, ⟨*W*⟩*/*10), and self-connections are not allowed. The corresponding percentage degree is set to *β* = 0.1. The leak term is set to *α* = 0.01 mHz. External currents of intensity *RI* = *RI*_1_ = · · · = *RI*_*K*_ = 1 mV are delivered to randomly selected neurons every *τ* = 0.01 ms over a total of 20,000 simulation time steps (equivalent to 2 seconds of activity). The same hyperparameter configuration is used to generate the data in Panel A of Fig. 1.

For each average synaptic weight, the total number of spikes is computed to examine the SNN’s global activity. The median and inter-quartile range across 8 different experimental repetitions are reported. In parallel, the information-based complexity is evaluated for the same set of hyperparameters to provide insight into the information richness of the spiking patterns. Specifically, the complexity is defined as the length of the LZW dictionary, i.e., the unique patterns identified by the algorithm applied to the time series composed of the concatenation of the activity of all the neurons. The median and inter-quartile range of the complexity across the same number of experimental repetitions as above are reported. To quantify the distance between the peak of the LZW complexity and the critical point identified in our analysis, we perform a pseudo-grid search by computing the absolute relative error for 30 additional hyperparameter configurations, either near or far from the boundaries of the three validity conditions: *ατ*_ref_ *<* 1*/*2, *θτ N/*(*RIτ*_ref_) ≥ 2, and *θτ Nα/*(*RI*) *<* 1. This enables us to effectively monitor potential anomalous behaviors and gain further insights.

### Self-organized quasi criticality

To present a computational example of self-organized quasi-criticality, we implement a local learning rule for synaptic weights, modeled as an iterative process that adjusts the network’s synthetic dynamics to approach a heuristic state in which half of the neurons are firing. The update of the average synaptic weight, occurring every *τ*_ref_, follows the equation:

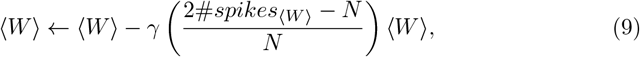

where the product between *γ* = 0.1 and #*spikes*_⟨*W* ⟩_ determines the step in terms of the percentage of the previous average synaptic weight (with initial ⟨*W*⟩ = 0.5). This framework provides an approach to analyzing how, under the model’s assumptions, synaptic weights evolve by assessing the network’s alignment (in terms of absolute relative error) with theoretical predictions. The numerical experiment examines this error as a function of the external input *RI* (in the range 2-8 mv) and the threshold *θ* (within the interval 10-200 mv). The remaining hyperparameters are fixed as following: *N* = 1000, *τ* = 0.1, *α* = 10^−4^, *τ*_ref_ = 1.

## Acknowledgements

We would like to thank Dr. Simone Russo for the useful discussions.

## Contributions

R.F. and A.B. conceived the study and handled the mathematical proofs. R.F and F.C. developed the code. R.F., F.C., and A.B. performed the synthetic analysis. R.F., F.C., L.M., and A.B. critically reviewed the manuscript. All the authors interpreted the results.

